# Methicillin-Resistant *Staphylococcus aureus* has Phenotypic Variation in *mecA* Expression that Alters Antibiotic Sensitivity

**DOI:** 10.1101/2025.05.28.656574

**Authors:** Dongzhu Ma, Rekha Arya, Beth Ann Knapick, Nadine Sadaka, Jewelia Rempuszewski, Claudette Moul, Jonathan B. Mandell, Neel B Shah, James B. Doub, Charlie Gish, Dana Parker, Stefanie A. Sydlik, Hunter Wood, Vaughn S. Cooper, Anthony R. Richardson, Robert M.Q. Shanks, Kenneth L. Urish

## Abstract

Methicillin resistant *Staphylococcus aureus* (MRSA) sepsis has a high rate of morbidity and mortality. Multiple clinical studies have demonstrated improved outcomes when MRSA sepsis is treated with dual antibiotic therapy that includes a β-lactam antibiotic such as cefazolin. This is a paradox as MRSA should be inherently resistant to this class of antibiotics. We report a serendipitous observation revealed a phenotype where MRSA became sensitive to cefazolin when cultured in a physiologic relevant media of fetal bovine serum as well as in synovial fluid. This could be observed across multiple clinical isolates. Expected resistance was maintained when cultured in Muller Hinton Broth (MHB). MRSA β-lactam antibiotic resistance is mediated by PBP2a, a penicillin-binding protein encoded by *mecA*. We hypothesized that this phenotype of antibiotic sensitivity in physiologic medium was based, in part, on levels of PBP2a expression and post-translational modifications of peptidoglycan wall teichoic acid (WTA). We therefore conducted quantitative RT-PCR analysis and Western blotting which demonstrated limited *mecA* expression in the mRNA level and limited PBP2a protein level when cultured in FBS or synovial fluid as compared to the clinical microbiology standard MHB, respectively. Whole genome sequencing of loss of function mutants generated through serial passaging in FBS revealed that the *clp* family of proteins and *rpo* genes were involved in β-lactam resistance. Cell wall peptidoglycan analysis suggested that WTA glycosylation was altered between β-lactam resistant and sensitive MRSA phenotypes. Together, this suggests pathways for *clpP*, *rpoB,* and WTA glycosylation can be new potential targets for MRSA treatment.

**IMPORTANCE:** This study demonstrates that MRSA has phenotypic variation in *mecA* expression, PBP2a protein levels, and patterns of wall teichoic acid glycosylation based on environmental conditions which can result in a phenotype of β-lactam antibiotic sensitivity. The presence of an antibiotic resistance gene does not necessarily result in antibiotic resistance. This provides a possible explanation for the clinical observations of potential therapeutic benefit of β-lactam antibiotics in dual therapy treatment of MRSA sepsis and suggests new therapeutic strategies to combat multidrug resistant bacteria. More importantly, identifying genes potentially responsible for this phenotype offers new potential therapeutic targets for the treatment of multidrug resistant infection.

## Introduction

*Staphylococcus aureus* bacteremia has a high morbidity and mortality (1), but not all strains of *S. aureus* are the same. Outcomes and treatment are dependent on the antibiotic resistance profile in which methicillin-resistant *Staphylococcus aureus* (MRSA) bacteremia can have twice the mortality rate of methicillin-sensitive *S. aureus* (MSSA) bacteremia (2). The difference in antibiotic resistance between MRSA and MSSA to β-lactam antibiotics is conferred from the presence of the *mecA* gene (3, 4). It encodes a penicillin-binding protein (PBP2a) with low affinity for these antibiotics. Clinically, this resistance is determined primarily through Clinical and Laboratory Standards Institute (CLSI) microdilution assay using Mueller-Hinton Broth (MHB) (5). The class of antibiotic used to treat *S. aureus* infection is dependent on the results of this assay. Infectious Diseases Society of America (IDSA) guidelines recommend cefazolin, a first-generation cephalosporin as primary treatment for MSSA, and either daptomycin or vancomycin for MRSA (6).

However, multiple clinical studies have demonstrated possible therapeutic benefits and improved clinical outcomes in MRSA sepsis when vancomycin or daptomycin was combined with cefazolin, when compared to vancomycin or daptomycin monotherapy alone. A prospective randomized clinical study comparing dual therapy to standard of care monotherapy was halted as the monotherapy group had higher rates of mortality as compared to combination therapy (8). In a second randomized prospective study, dual therapy decreased the overall time of bacteremia as compared to monotherapy (9). The presence of an antibiotic resistance gene does not guarantee its expression and there can be variation between an antibiotic resistance genotype and phenotype induced by different environmental conditions (7,10). Therefore, quantifying clinical antibiotic sensitivity using the CLSI protocol may fail to replicate the actual metabolic and environmental conditions of the infection leading to a variation between the actual and measured antibiotic resistance phenotype (11, 12). Based on this and the unexpected benefit of dual therapy in MRSA sepsis, we questioned if there was variability in the resistance phenotype of MRSA with the β-lactam cefazolin between physiological relevant media and MHB, the media used in the CLSI protocol. We hypothesized that culturing MRSA in a physiologic relevant media would result in a sensitivity to cefazolin as compared to culturing in MHB. We demonstrated that MRSA cultured in MHB was resistant to β-lactam antibiotics but became sensitive in physiologic relevant mediums of serum or synovial fluid. This phenotypic sensitivity was associated with a decrease in expression of PBP2a and post-translational modification of wall teichoic acid glycosylation.

## Results

### MRSA strains become susceptible to beta-lactam antibiotic cefazolin when grown in FBS

In a serendipitous observation, our group observed MRSA cefazolin resistance was a phenotype that was dependent on culture conditions. Resistance to cefazolin, a first-generation cephalosporin, as determined by MIC, was variable based on media type. MRSA laboratory strain (JE2) and 9 MRSA clinical isolates were resistant to cefazolin in MHB (greater than or equal to 8 µg/mL) (**Figure 1A**), the culture media used in clinical microbiology laboratories. However, these isolates became sensitive (MIC<=2; **Figure 1B**) to cefazolin in a culture media representing conditions similar to physiologic serum (fetal bovine serum, FBS). These observations are supported by the counterintuitive results of clinical studies that have demonstrated therapeutic benefit from dual antibiotic therapy with cefazolin in MRSA sepsis (8, 9).

**Figure 1.**
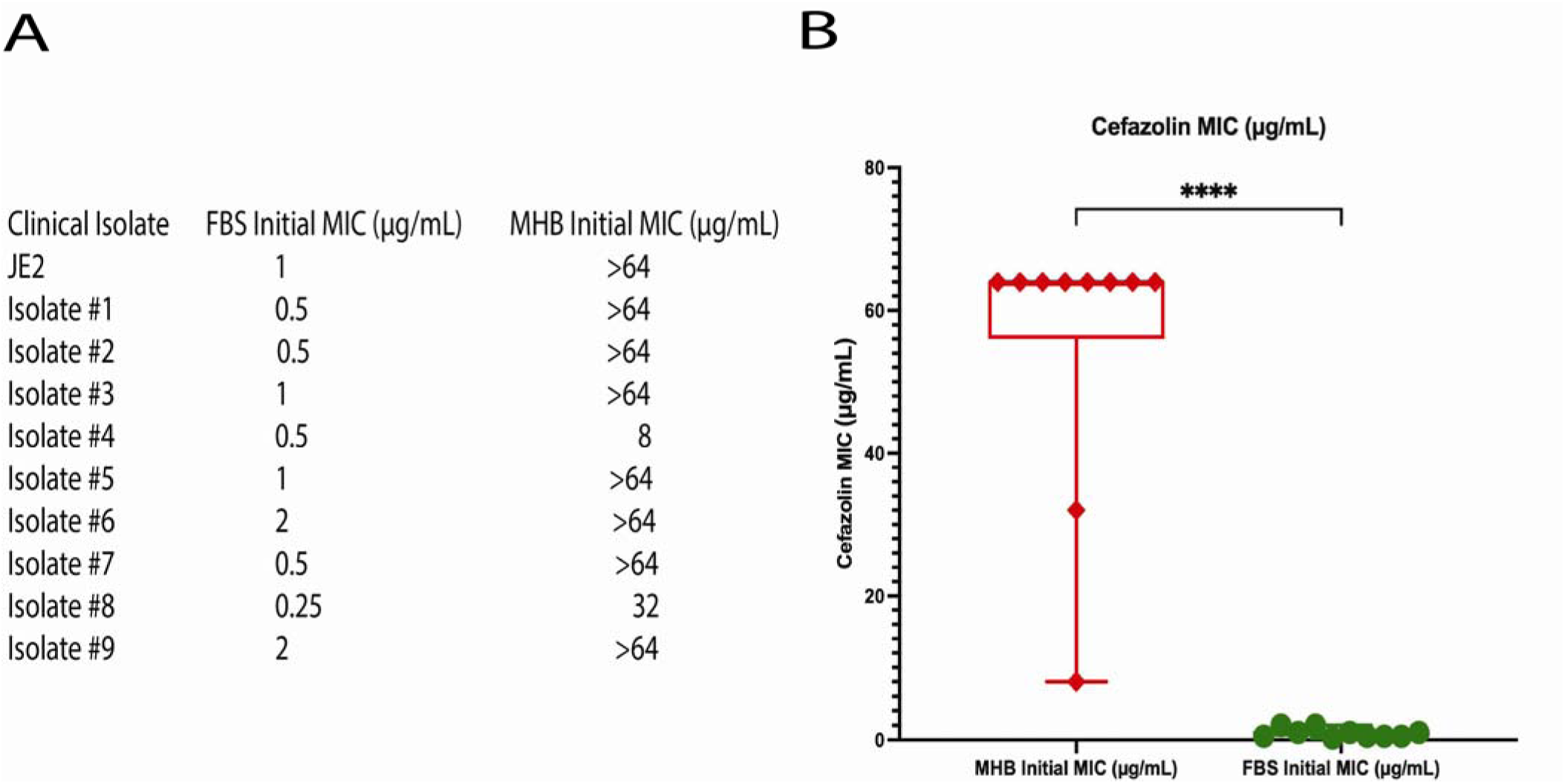
MRSA becomes susceptible to the beta-lactam antibiotic cefazolin when grown in FBS. (A) Cefazolin MIC assays were performed on JE2 and 9 clinical MRSA isolates in Mueller Hinton Broth (MHB) and fetal bovine serum (FBS), respectively. (B) MRSA isolates cultured in FBS were sensitive in cefazolin as compared to MRSA isolates cultured in MHB (****p<0.0001).

### MRSA isolates had decreased mecA mRNA in physiologic media that correlated with cefazolin sensitivity

To determine if this was observed in other physiologic environments, the experiment was repeated in an additional type of physiologic media representative synovial fluid. Serum is not a defined media, but a defined version of synovial fluid termed pseudo synovial fluid (pSF) has been validated (13). This pSF is designed to replicate key characteristics of natural synovial fluid. When these same clinical isolates were cultured in pSF, the cefazolin MIC remained relatively low at 4 µg/mL.

To determine a mechanism for this phenotype that was observed in both serum and synovial fluid environments, we first wanted to determine if this was before or after translation of the primary gene associated with β-lactam resistance, *mecA*. We hypothesized that the phenotypic sensitivity to cefazolin was based on variable expression of the *mecA* gene. To test this hypothesis, we measured the expression levels of *mecA* (mRNA) and PBP2a, the protein associated with the *mecA* gene, with quantitative PCR and Western blot, respectively, in different growth medium (**Figure 2**). GAPDH was used as a reference control. *mecA* expression had a large and statistically significant reduction in FBS and pSF as compared to TSB and MHB (p<0.0001, **Figure 2A**). Protein levels of PBP2a were decreased in FBS and pSF media as compared in TSB and MHB determined by Western blot (**Figure 2B**). These finding suggested that *S. aureus* JE2 *mecA* expression decreased in physiologic media (FBS and pSF) as compared to standard laboratory media (MHB and TSB).

**Figure 2.**
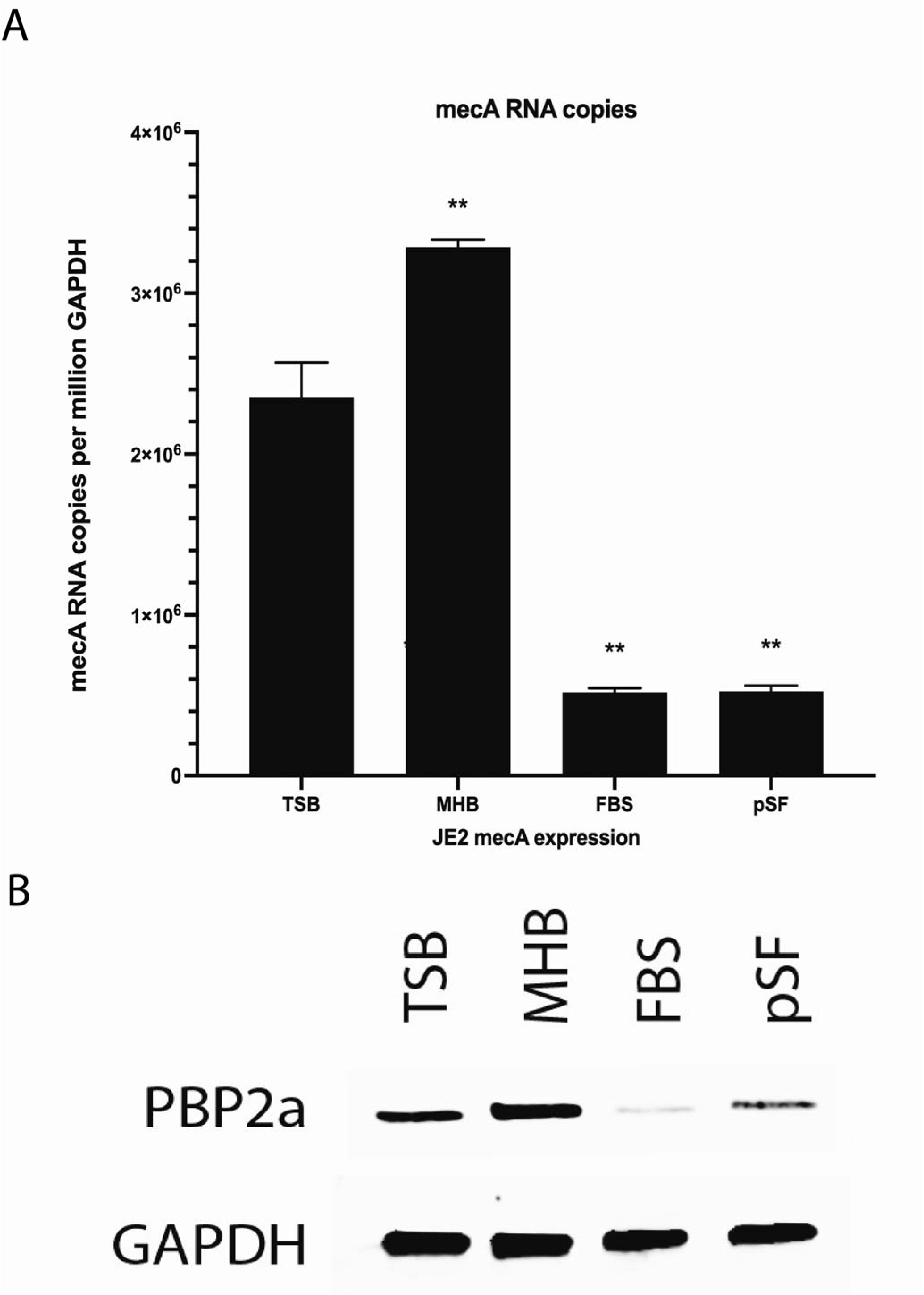
Physiologic culture media altered *mecA* mRNA and PBP2a expression. Quantitative real-time PCR and Western blotting were performed on JE2 cultured in TSB, MHB, FBS, and Pseudo synovial fluid (pSF). (A) MRSA (JE2) had higher levels of *mecA* mRNA when cultured in TSB and MHB as compared to FBS and pSF at a 4-hour time point (****p<0.0001). (B). MRSA (JE2) had higher levels of PBP2a when cultured in TSB and MHB as compared to FBS and pSF at a 4-hour time point.

### JE2 resistant to cefazolin in physiologic media maybe driven by clp and rpo genes

To identify possible genes responsible for this phenotypic cefazolin sensitivity, MRSA (JE2) was serially passaged in FBS until the β-lactam sensitivity phenotype was lost. We obtained 12 isolates that were initially sensitive to cefazolin in physiologic medium, lost the phenotype, and developed comparable resistance to cefazolin in physiologic and laboratory media **(Figure 3A)**. Similar to the changes observed between culturing MRSA in physiologic or laboratory media, loss of β-lactam sensitivity in these mutants was associated with higher expression levels of *mecA* when compared to the wild-type strain (**Figure 3B**). Additionally, we did not observe any notable changes in PBP2a levels in resistant mutants when compared with the sensitive wild type (**Figure 3C).** Based on these findings, we postulated that this lack of changes may be attributed to alternative mechanisms affecting PBP2a distribution, modification, degradation, or conformational changes or other pathways.

**Figure 3.**
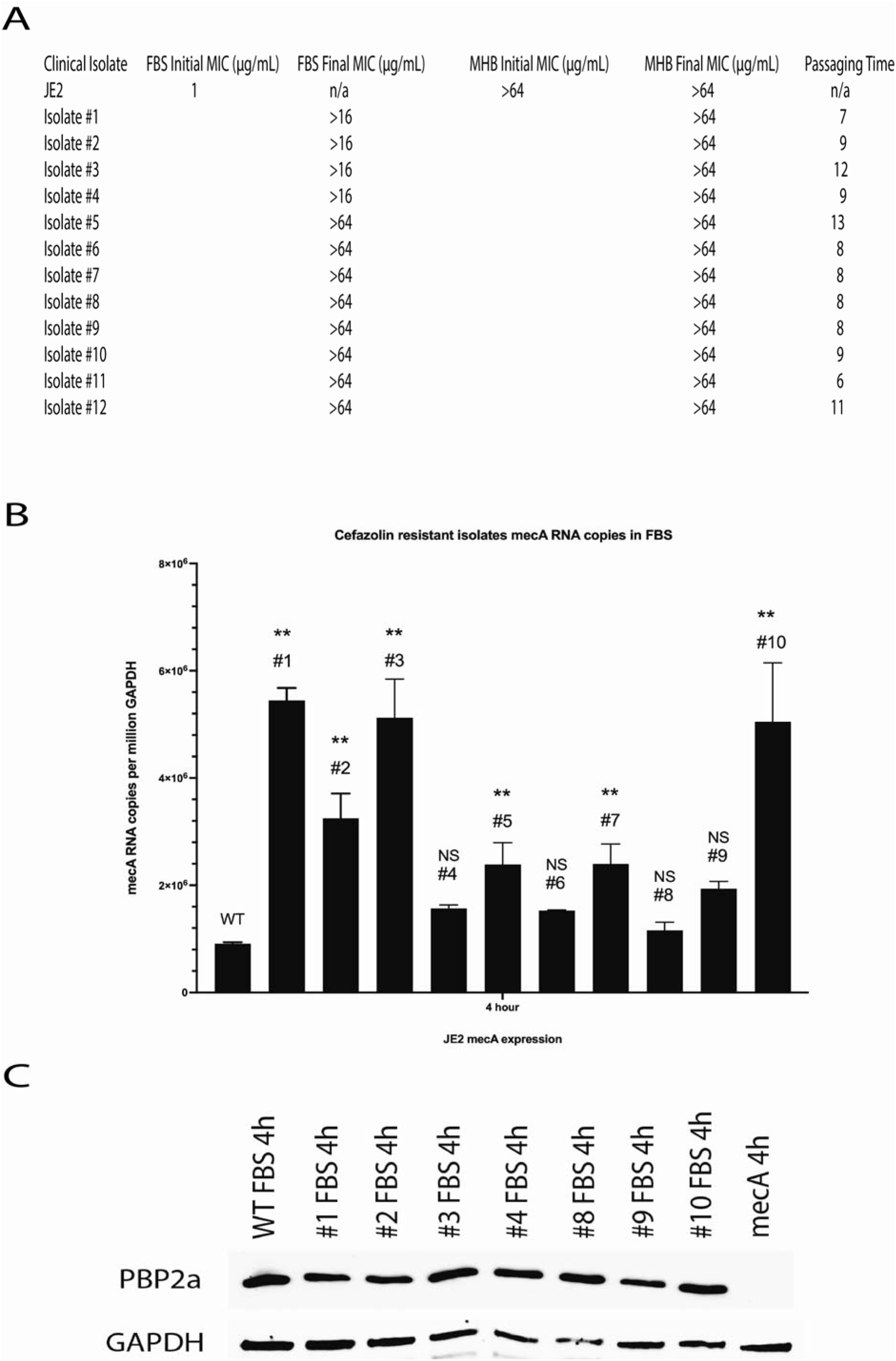
Serially passaged MRSA that has lost the phenotype of beta-lactam sensitivity in physiologic media express more *mecA* mRNA than that of wild type in FBS. (A) JE2 was serially passaged in FBS with cefazolin until resistance developed and was comparable to MHB MIC. (B) Quantitative real-time PCR of these MRSA cefazolin resistant isolates in FBS was performed. These mutants expressed higher levels of *mecA* mRNA cultured in FBS compared with wild type at a 4-hour time point (NS=no significance, **p<0.01). (C) Western blotting of these MRSA cefazolin resistant isolates in FBS was performed. No appreciable differences in PBP2a levels between wild type and mutants were observed.

Whole genome sequencing (WGS) was completed to identify the mutations that were associated with the loss of the phenotype (**Table 1**). Our genomic analysis revealed distinct mutations in either *rpo* or *clp* genes in serially passaged loss of function mutants. For the *rpo* family, there were 7 isolates with missense mutations in the *ropB* gene (A232E, S385I, S385R, E662V, A742E, S964Y, and V1144D), one isolate with a missense mutation in the *rpoC* gene (A6D), and 2 isolates with a missense mutation in the *rpoD* gene (G320C, L321P). For the *clp* family, 2 isolates had missense mutations in the *clpP* gene (D27Y, R28C), 2 isolates had a missense mutation in the *clpX* gene (K83N, L335P), and one isolate had a clpX frame shift mutation (-1bp). Mutations in *rpoB/rpoC*, encoding the β and β′ subunits of the RNA polymerase, can facilitate adaptation to diverse environmental, antibiotic stresses, and are responsible for resistance phenotypes (14–16). *rpoB* and *rpoC* have been reported to be involved in bacteria β-lactam resistance (16–18).

**Table 1.**
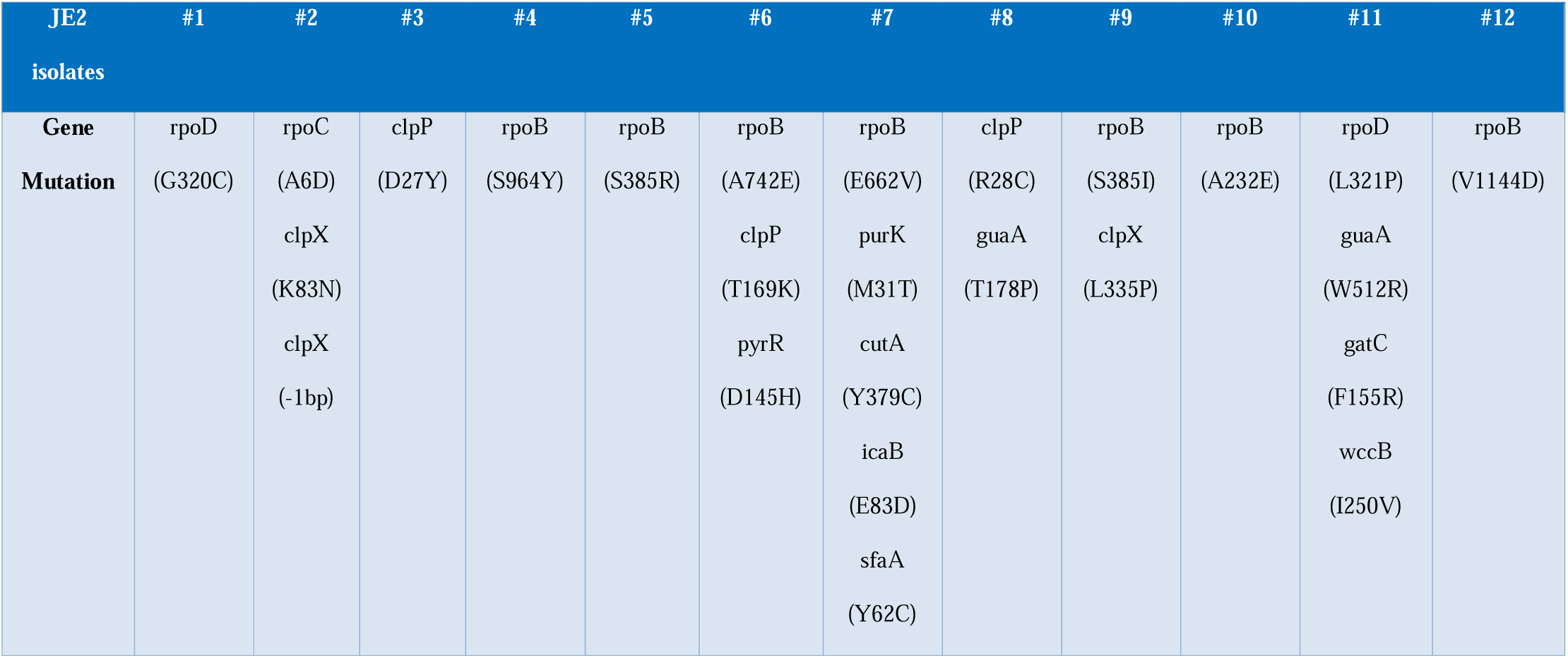
NGS genomic DNA analysis of cefazolin resistant isolates.

Based on the above, we hypothesized that these mutations in *rpo* or *clp* may play a role in cefazolin resistance. To test this hypothesis, we measured the MIC of available mutant strains of *clp, clpB, clpC, clpP, rpoE, and rpoF* from the NARSA library. Importantly, we verified the accuracy of these transposon mutants with PCR. Comparable to the *clpP* loss of function mutant from the serial passaged experiments (isolate #3), the *clpP* transposon mutant from the NARSA library was resistant to cefazolin in FBS, which was the opposite of the phenotype of the wild type control strain. The other *clp* transposon mutants had a comparable phenotype to the wild type where there were sensitive to cefazolin in FBS. This suggests that *clpP* is involved in β-lactam resistance in FBS. (**Table 2**).

**Table 2.** Cefazolin MIC assays of NARSA strains (μg/mL)

| Strains | WT (JE2) | clp | clpB | clpC | clpP | Resistant isolate #3 (clpP) | rpoE | rpoF |
| --- | --- | --- | --- | --- | --- | --- | --- | --- |
| **TSB** | 8 | 8 | 8 | 8 | 32 | ≥ 64 | 8 | 2 |
| **MHB** | ≥ 64 | ≥ 64 | ≥ 64 | ≥ 64 | ≥ 64 | ≥ 64 | 32 | 4 |
| **FBS** | 1 | 1 | 1 | 1 | 8 | 32 | 2 | 1 |

In our genomic analysis, the majority of mutations occurred in the *rpo* family of genes, and so we attempted to understand the role of *rpoB* in mediating cefazolin resistance in physiologic media. RpoB protein levels were measured in each of the serially passaged isolates and it was noted that RpoB protein levels were variable in each of the isolates suggesting that expression level may not be associated with this phenotype of beta-lactam resistance in physiologic media (Figure 4A). Using the NARSA library available *rpo* and *clp* mutations (*clp, clpB, clpC, clpP, rpoE, rpoF*, and *mecA*), we then determined if these genes would alter *rpoB* expression. We did not observe any significant difference in the protein levels of *rpoB* from these NARSA library *clp* and *rpo* mutations (**Figure 4B**). Finally, to test if the loss of *rpoB* could lead to β-lactam resistance, we attempted to create a *rpoB* gene deletion mutant by homologous double crossover recombination using pKFT-rpoB vector (19) (**Supplemental data**). This was attempted more than four times and was not successful. The failure to generate a viable *rpoB* deletion mutant demonstrates that *rpoB* is essential for the survival of *S. aureus* JE2. Overall, these findings suggested that the rpoB protein level was not associated with cefazolin resistance and was more likely attributed to an alternative mechanism such as conformational or functional change.

**Figure 4.**
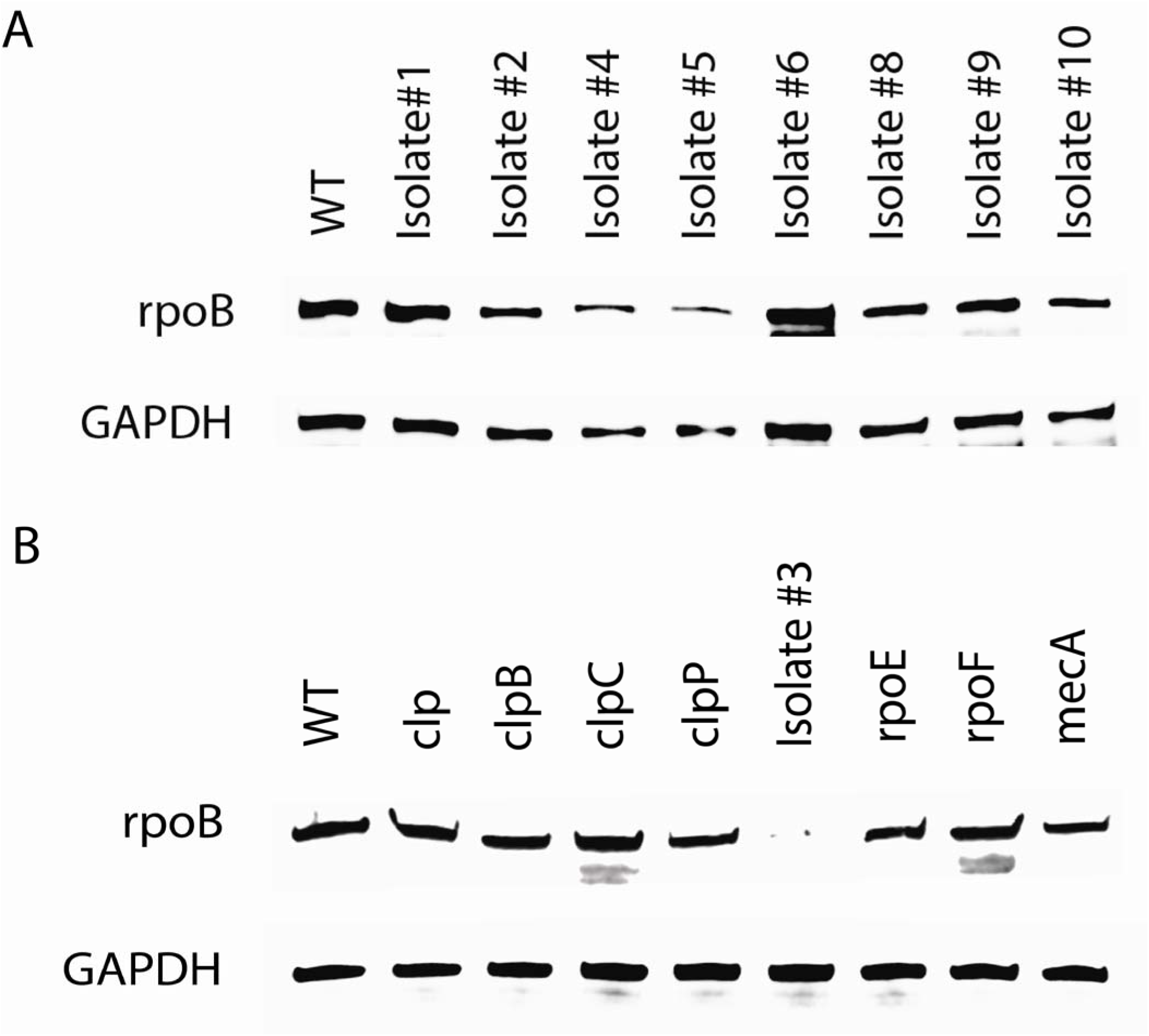
NARSA library *clp* and *rpo* mutants had comparable levels of rpoB protein. JE2 *rpoB* expression was measured by Western blotting in different strains. (A) rpoB protein levels of JE2 cefazolin resistant isolates in TSB. (B) rpoB protein levels of mutant strains from NARSA Library in TSB.

#### Glycosylation of cell wall teichoic acids (WTA) is altered in FBS as compared to MHB

Cell wall teichoic acids (WTA) play a crucial role in the structural integrity and pathogenicity of *S. aureus*. Different glycosylation patterns of WTA have been demonstrated to alter β-lactam sensitivity in MRSA (20, 21). Further, environmental conditions, such as the nutrient composition of the growth medium, can affect WTA modifications. As a result, we hypothesized that WTA glycosylation patterns may be altered in physiologic growth medium as compared to standard laboratory medium. To assess this, we compared WTA glycosylation patterns in JE2 cultured in MHB and FBS, using the ratio of peaks at 5.1 ppm and 4.7 ppm corresponding to the α (1–4) GlcNAc proton of WTA and the β (1–3) GlcNAc proton of WTA, respectively. After extracting WTA (20, 22–24), proton nuclear magnetic resonance (NMR) spectra demonstrated a decrease in α (1–4) GlcNAc to β (1–3) GlcNAc ratio from 0.0943 to 0.0581, confirming changes in the glycosylation pattern when cultured in MHB versus FBS (**Figure 5**). Overall, these results concluded that the glycosylation pattern of WTA contributes to increased JE2 cefazolin susceptibility in FBS.

**Figure 5.**
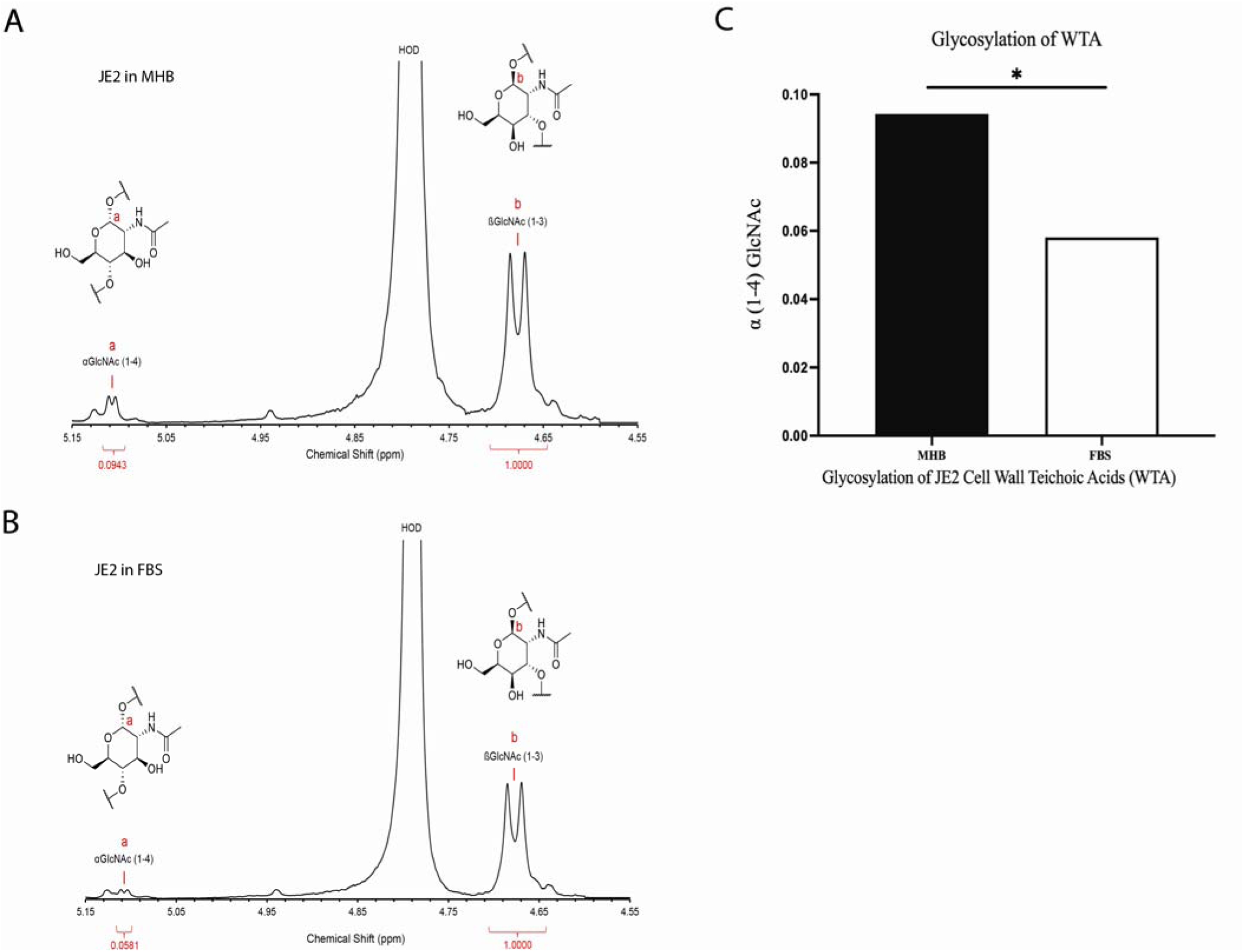
Glycosylation of JE2 Cell Wall Teichoic Acids (WTA) decreased in FBS than that in MHB. (B). Glycosylation of WTA of JE2 in MHB. α (1–4) GlcNAc of WTA is 0.0943, β (1–3) GlcNAc of WTA is 1.0). (B) Glycosylation of WTA of JE2 in FBS. α (1–4) GlcNAc of WTA is 0.0581, β (1–3) GlcNAc of WTA is 1.0). (C). Compare of JE2 WAT α (1–4) GlcNAc in MHB and FBS.

## Discussion

Methicillin-resistant *Staphylococcus aureus* (MRSA) infections can have high morbidity and mortality. The resistance to β-lactam antibiotics is determined by the *mecA* gene which encodes the penicillin-binding protein PBP2a. Penicillin-binding proteins (PBP) are a group of proteins necessary for cell wall synthesis where, as the name implies, penicillin has a natural binding affinity. By binding to PBP, beta-lactam antibiotics inhibit bacterial cell wall biosynthesis leading to cell lysis and death (25). MRSA developed a resistance to β-lactam antibiotics via the *mecA* gene which encodes an alternate penicillin-binding protein, PBP2a. This prevents β-lactam binding and allows cell wall synthesis to continue.

Here, we observed MRSA antibiotic resistance to β-lactam antibiotics can be variable. We observed that MRSA resistance to β-lactam was altered when cultured in physiologic medium (FBS or synovial fluid) as compared to traditional laboratory medium (TSB or MHB). In physiologic culture medium, MRSA clinical isolates developed a phenotype that was sensitive to β-lactam antibiotics and reverted when cultured in MHB, the media used to clinically determine resistance. This sensitivity was based, in part, on limited *mecA* expression levels, decreased PBP2a protein levels, and WTA post-translational modifications.

Based on the characteristics of PBPs in antibiotic resistance, we assessed cefazolin resistance by measuring *mecA* expression and PBP2a level. Compared to cultures in MHB, *mecA* mRNA and PBP2a expression levels were reduced in FBS (**Figure 2**). We hypothesized that environmental factors present in host conditions of *S. aureus* infections could influence *mecA* expression thereby impacting antibiotic efficacy. Pseudo synovial fluid (pSF) was used to mimic the *S. aureus* host joint microenvironments.

Our results demonstrated that antibiotic susceptibility could vary depending on the culture medium. Specifically, JE2 exhibited increased susceptibility to cefazolin in pSF and FBS, compared to standard laboratory media MHB. This was a result of decreased *mecA* expression and subsequent levels of PBP2a (**Figure 2**).

To determine a mechanism driving the change in resistance, WGS identified that rpoB played an important role in the phenotype. RpoB is also a target for rifampicin, and that site mutation of *rpoB* can confer rifampicin resistance (26–29). The *rpoB* gene encodes the β subunit of bacterial RNA polymerase. Rifampicin (RMP) binds within the RNAP β subunit (30). Mutations in *rpoB* can offer resistance to rifampicin because they alter the drug-binding residues of the protein, thereby reducing affinity for these antibiotics (26, 30, 31). Further studies showed that the *rpoB* mutation resulted in decreased expression levels of anaerobic respiration and fermentation genes, while 11 genes such as *mecA* were specifically upregulated (16). This may connect rpoB with β-lactam antibiotic resistance. When bacteria are placed under diverse selective pressures, they often adapt rapidly by mutating to the most conserved positions in *rpoB* (15). In this study, JE2 cefazolin resistant isolates in FBS were obtained by serial passages. Genomic DNA analysis of these isolates revealed that *rpoB* was most likely involved in the loss of phenotype and developing resistance to cefazolin. However, the exact mechanism of resistance remains unknown. Changes in rpoB conformation or other mechanisms may be involved, such as reduced cell wall permeability, cell wall thickness, or enhanced efflux pumps.

Peptidoglycan (PG) is the major structural component of the bacterial cell wall. Gram-positive bacteria are characterized by a thick cell wall composed of a peptidoglycan matrix embedded in anionic polyol phosphate polymers, known as teichoic acids (32–34). WTA is a major component of the *S. aureus* cell wall and plays an important role in the bacteria’s physiology (adhesion, colonization, and cell division) (35), virulence, biofilm formation, and resistance to antimicrobial molecules (36, 37). WTAs covalently attached to peptidoglycan via a phosphodiester linkage (38). WTA glycosylation is influenced by environmental conditions and contributes to resistance to antibiotics (20). Glycosyltransferases TarS and TarM are involved in WTA α-O-GlcNAcylation and β-O-GlcNAcylation, respectively, and are essential for normal cell function (20). We observed that culturing JE2 in a physiologic media altered its WTA glycosylation (α (1–4) GlcNAc). This suggests that WTA post-translational processes may also play a role in this observed β-lactam resensitization.

While these findings provide a mechanistic underpinning for the potential benefit of dual therapy for MRSA bacteremia, there are limitations in this study. Only 9 clinical MRSA strains and 12 JE2 cefazolin-resistant isolates were analyzed, as well the number of isolates containing certain types of *rpoB* mutations was still limited. Because *rpoB* is a gene essential for bacterial survival, manipulation of the *rpoB* gene can easily lead to lethal consequences. Therefore, it was difficult to conduct site-directed mutagenesis studies on *rpoB* genome manipulation.

## Conclusion

Expression of *mecA* and PBP2a has been extensively studied in relation to methicillin (oxacillin) resistance. We observed that MRSA could become sensitive to β-lactam antibiotics (cefazolin) when cultured in a physiologically relevant media. This phenotype was associated with higher *mecA* expression and PBP2a protein levels. *rpoB* and *clpP* genes were associated with this phenotype. This identifies a potential new therapeutic targets for MRSA infections and suggests a new strategy for treatment of other multidrug resistant infections.

## Materials and Methods

### Bacterial strains, plasmids and growth conditions

The bacterial USA300 FPR3757 strain (BAA-1556, JE2) was purchased from The American Type Culture Collection (ATCC). NR-48501 *Staphylococcus aureus* subsp. aureus Nebraska Transposon Mutant Library (NARSA) was obtained from BEI resources (https://www.beiresources.org/Home.aspx). Trypticase soy broth (TSB), Mueller–Hinton II broth (MHB) were purchased from BD (USA) and sterilized by autoclaving. Fetal bovine serum (FBS) was purchased from Invitrogen (Thermo Fisher Scientific, USA). *Staphylococcus aureus* strains were growth in TSB medium with or without appropriate antibiotics. Ampicillin, Erythromycin, Tetracycline, Chloramphenicol, cefazolin, Vancomycin, Rifampin, Lysostaphin, Anti-rpoB and Anti-PBP2a of MRSA were purchased from Sigma-Aldrich (USA). Anti-GAPDH (GA1R) antibody and goat anti-mouse IgG (H+L) secondary antibody-HRP were purchased from Invitrogen (Thermo Fisher Scientific, USA). SDS-PAGE and protein electrophoresis system and ECL reagents were purchase from Bio-Rad (USA). iBlot3 protein semi-dry transfer system was purchased from Invitrogen (Thermo Fisher Scientific, USA).

### Minimum inhibitory concentration (MIC) assay

The minimum inhibitory concentration (MIC) of cefazolin, Vancomycin and Rifampin for *S. aureus* JE2 in suspension was determined using CLSI assay protocols, incubating freshly plated cultures at 0.5 × 10^6^ CFU/mL for 24 hours with serial dilutions of each antimicrobial and observing inhibition of bacterial growth based on turbidity or using the PrestoBlue viability assay (Thermo Fisher Scientific, USA) on the treated cultures according to the manufacturer’s instructions and read on a SynTek microplate reader measuring absorbance at 600 nm. Antibiotics concentrations ranged 0, 0.25, 0.5, 1, 2, 4, 8, 16, 32, 64 µg/mL.

### Pseudo synovial fluid (pSF) preparation

Pseudo synovial fluid (pSF) was prepared by combining 3 mg/mL Hyaluronic Acid (HA), 9 mg/mL Fibrinogen (Fg), and 10 mg/mL Albumin (Alb) in Dulbecco’s Modified Eagle Medium supplemented with 5% Non-Essential Amino Acids (DMEM) (13). Each component was also tested individually at the stated concentration in DMEM. The minimum inhibitory concentration (MIC) of JE2 was assessed described as above.

### Serial passage and cefazolin resistant isolates selection

Using a well-characterized MRSA strain, USA300 FPR3757 strain (BAA-1556, ATCC), we selected for cefazolin resistance by serial passage. A well-isolated colony was picked from an overnight agar plate and inoculated into a TSB medium to achieve the specified turbidity by comparing to a 0.5x McFarland turbidity standard (1×10^8^/mL). Approximately 1×10^6^ cells were added to each well of a 96-well plate that was filled with FBS containing half diluted cefazolin. The plates were incubated at 37□C for overnight. We recorded antibiotic concentration where bacteria growth was inhibited (MIC). The FBS wells right below the MIC row were mixed and then pipetted into a 2 mL centrifuge tube. The MIC assay was repeated until the FBS MIC reached ≥64 μg/mL. Next ≥64 μg/mL well was transferred into sterile tube and mixed. Using an inoculating loop, a streak plate was performed and then incubated at 37□C for overnight. The streak plate was passaged for 10 days. After 10 days, culture clones overnight in 4 mL of TSB at 37□C. Repeat MIC assay with FBS. If the plate reaches MIC >16 μg/mL. The isolates are banked and then stored at -80□C.

### Genomic bacterial DNA isolation

Using MasterPure™ Gram Positive DNA Purification Kit (Lucigen Corp., USA) to isolate genomic DNA from *Staphylococcus aureus* samples. Clinical *Staphylococcus aureus* stock samples stored at -80 were streaked out on TSA plate and then incubated overnight at 37□C. A single colony from TSA plate was picked then inoculated in 4 mL TSB medium and grown overnight at 37□C in a 250 rpm shaker. 1.5 mL of the overnight grown bacterial culture was pelleted. 150 μL of TE Buffer was added and then vortexed to resuspend the cell pellet. 1 μL of Ready-Lyse Lysozyme and 5 μL of lysostaphin (5 mg/mL) were added to each resuspended pellet of bacteria. The samples were incubated at 37□C until bacterial cell wall is destroyed. 150 μL of Proteinase K/Gram Positive Lysis Solution was added to the sample and then mixed thoroughly. Samples were incubated at 65□C for 15 minutes, vortexing briefly every 5 minutes. The samples are cooled to 37□C. Samples were placed on ice for 5 minutes and then 175 μL of MPC Protein Precipitation Reagent was added to 300 μL of lysed sample and then vortexed vigorously for 10 seconds. The debris was pelleted by centrifugation at 4□C for 10 minutes at 12,000 x g. The supernatant was transferred to a clean microcentrifuge tube and the pellet was discarded. 1 μL of RNase A (5 μg/μL) was added to each sample and mixed thoroughly. Samples were then incubated at 37□C for 10 minutes. 500 μL of isopropanol was added to the recovered supernatant. The tubes were inverted 30-40 times. The DNA was pelleted by centrifugation at 4□C for 10 minutes at 12,000 x g. A pipet tip was used to remove the isopropanol without dislodging the DNA pellet. The pellet was rinsed with 70% ethanol. The DNA was resuspended in 100 μL of TE Buffer. The genomic DNA was used as template in following PCR reactions and sequencing.

### Bacterial whole genome sequencing

Next generation whole genome sequencing was performed by SeqCoast Genomics LLC. (https://seqcoast.com/)

### Preparation RNA Standards for JE2 *S. aureus mecA* gene

The standard curve method was selected to quantify *mecA* mRNA copies. To obtain RNA standards, the *mecA* gene fragments was amplified from JE2 *S. aureus* strain. The primers for *mecA* and GAPDH genes were JE2-mecA-standard-out-F (5′-TTTgaattcCCGTTCTCATATAGCTCATCATACA-3′), JE2-mecA-standard-out-R (5′-ATTggatccTGTTGGTCCCATTAACTCTGAA-3′). JE2-GAPDH-standard-out-F (5′-AGGTgaattcGAGGTAGTTGATGGTGGTTTC-3′), JE2-GAPDH-standard-out-R (5′-ATGggatccCGTTATCATACCAAGCTGCAAC-3′). The fragment was cloned into pBlueScript II SK+ (Agilent Technologies, USA) and linearized by *XbaI* as a DNA template for RNA in vitro transcription. Standard RNA template was produced by using Ambion® MEGAscript® kit (Thermo Fisher Scientific, USA) following kit manual instruction. Standard RNA concentration was determined by NanoDrop™ 2000 Spectrophotometers (Thermo Fisher Scientific, USA).

### Isolation of RNA and quantitative RT-PCR analysis

RNA isolation and quantitative reverse transcription polymerase chain reaction (RT-PCR) were performed by following the manufacturer’s instruction of the product. *S. aureus* was grown in 4 mL of TSB medium supplemented with appropriate antibiotics at 37□C for overnight (about 16 hours). 200 μL overnight growth was added into 4.0 mL fresh TSB medium and then grown at 37□C for 2 hours. All 4 mL of bacterial culture was pelleted. 100 μL of TE Buffer was added to resuspend the cell pellet. 5 ul Lysostaphin (5 mg/mL) was added to the resuspended pellet of bacteria. The samples were placed at 37□C for 15 minutes to destroy the cell wall. Total RNA was extracted by using TRIzol® Max™ Bacterial RNA Isolation Kit (Thermo Fisher Scientific) and reverse transcription of RNA to single-stranded cDNA was performed using SuperScript IV Reverse Transcriptase (Thermo Fisher Scientific, USA). The newly synthesized cDNA was used immediately or frozen at -80□C.

Quantitative RT-PCR analysis was performed by using the CFX96 Real-Time System (BioRad, USA) and TaqMan^TM^ Fast Advanced Master mix (Thermo Fisher Scientific, USA). The cycling conditions used were initial cycle 95 C for 2 min, followed by 45 cycles of 95 C for 10 sec and 60□C for 60 sec. The primers for *mecA* and *GAPDH* genes were JE2-mecA-qPCR-F (5′-TTGTAGCTAGCCATTCCTTTATCT-3′), JE2-mecA-qPCR-Probe (5′-ACCACCCAATTTGTCTGCCAGTTTC-3′), JE2-mecA-qPCR-R (5′-CCGGTACTGCAGAACTCAAA-3′). JE2-GAPDH-qPCR-F (5′-CACCAAGAGTTAGACGGTTCTG-3′), JE2-GAPDH-qPCR-Probe (5′-TGACTACAATTCACGCTTACACAGGTGA-3′), JE2-GAPDH-qPCR-R (5′-TGAGGTGCGTCTTGTGTATTT-3′). The actual RNA copies were calculated according to the standard curve. Statistical analysis was performed using a two-sided Student’s t-test or one-way ANOVA.

### Detection of PBP2a by Western blot

1:100 overnight grown *S. aureus* cells were inoculated in 4 mL growth medium for 2-4 hours. 1,500 μL *S. aureus* cells were collected by centrifugation at 5,000 x g for 10 min at 4 °C. The cell pellet was washed twice with 1.5 mL cold PBS and spun as before. The cell pellet was resuspended in 150 μL of TE buffer by vigorous vertexing. 5 μL of lysostaphin (5 mg/mL) was added to the cell resuspension. The resuspension was incubated at 37 °C for 15-30 min and the tube was inverted several times until the mixture became very viscous and transparent. The samples were sonicated on ice for 15 seconds. 150 μL of Lysis buffer (50 mM Tris-HCl, 7.5 pH, 1% Triton X-100, 20 μg/mL DNase, 10 μg/mL RNase, 1x protease inhibitor cocktail) was added to the mixture and then incubated on ice for 30 minutes. Centrifugation at 12,000 x g for 10 min at 4 °C. The supernatant is kept. The protein concentration of the lysates was measured using Bradford Assay (Bio-Rad) and samples were normalized to equal amounts of protein, 10 μg of total protein was separated using Mini-PROTEAN precast gels (Bio-Rad). Transfer the protein to PVDF membrane with iBlot™ 3 Western Blot Transfer system (Thermo Fisher Scientific). Blocking the PVDF membrane using EveryBlot Blocking Buffer (Bio-Rad) at room temperature for 2 hours. The primary antibody (1:2000 dilution in EveryBlot Blocking Buffer) of anti-PBP2a or GAPDH was applied to membranes overnight at cold room with agitation. A horseradish peroxidase (HRP) conjugated goat anti-mouse IgG secondary antibody (Thermo Fisher Scientific) (1:5000 dilution in EveryBlot Blocking Buffer) was incubated with the membrane for 2 hours. Clarity ECL Western blotting substrates (Bio-Rad) was used for visualization of proteins. The blot was obtained using ChemiDoc MP Systems (Bio-Rad) for chemiluminescent detection.

### Extraction, purification, and Analysis of Bacterial Wall Teichoic Acids (WTA)

Glycosylation of *Staphylococcus aureus* cell wall teichoic acid is influenced by environmental conditions and suggest that β-GlcNAc WTA may bring competitive advantage *in vivo* (20). WTAs were extracted and purified from the JE2 strains grown in 200 mL culture of MHB and FBS for 24 h at 37□C. Overnight cultures were centrifuged and washed with Buffer1 (50 mM MES, pH 6.5). The cultures were inactivated by treatment with phenol-ethanol (1:1 v/v) to a final concentration of 2%. The cells were collected by centrifugation at 2500g for 1 hour at 4□C and suspended in 0.05 M Tris, 2 mM MgSO_4_, pH 7.5 (0.5g wet weight/mL). The cell suspensions were incubated with lysostaphin (50 μg/mL) at 37□C for 3 hours with continuous stirring. MgCl_2_ and Benzonase were subsequently added to a final concentration of 1 mM and 50 UI/mL, respectively, and incubated at 37□C for 4 hours. The final concentration of Tris buffer was adjusted to 50 mM, and CaCl_2_ and Pronase added to a final concentration of 1 mM and 0.5 mg/mL, respectively. Samples were incubated for 16 hours at 37□C. The remaining insoluble cell debris were removed by centrifugation at 8,000g for 30 min. The supernatants were precipitated with 25% ethanol in presence of 10 mM CaCl_2_ and stirred for 16 hours at 4□C. The precipitates were removed by centrifugation at 8,000g for 30 min. The supernatants containing WTAs were precipitated with 75% ethanol in presence of 10 mM CaCl_2_ and stirred for 4 hours at 4□C. Then, WTAs were collected by centrifugation at 8,000g for 30 min and dissolved in water. The samples were dialyzed extensively against water at room temperature. After dialysis for 24-48 hours, the residues were lyophilized, and the solids were frozen and kept in -80□C freezer. The breakdown of glycosylation states of the WTA of each sample was analyzed by Dr. Sydlik and Dr. Hunter Wood from Carnegie Mellon University.

## Acknowledgements

Dr. Kenneth Urish is supported in part by the National Institute of Arthritis and Musculoskeletal and Skin Diseases (NIAMS R01 AR082167 and R03 AR077602).

## References

1. Turnidge JD, Kotsanas D, Munckhof W, Roberts S, Bennett CM, Nimmo GR, Coombs GW, Murray RJ, Howden B, Johnson PD, Dowling K, Australia New Zealand Cooperative on Outcomes in Staphylococcal S. 2009. Staphylococcus aureus bacteraemia: a major cause of mortality in Australia and New Zealand. Med J Aust 191:368–73.

2. Whitby M, McLaws ML, Berry G. 2001. Risk of death from methicillin-resistant Staphylococcus aureus bacteraemia: a meta-analysis. Med J Aust 175:264–7.

3. Fishovitz J, Hermoso JA, Chang M, Mobashery S. 2014. Penicillin-binding protein 2a of methicillin-resistant Staphylococcus aureus. IUBMB Life 66:572–7.

4. Peacock SJ, Paterson GK. 2015. Mechanisms of Methicillin Resistance in Staphylococcus aureus. Annu Rev Biochem 84:577–601.

5. Anonymous. 2015. M100-Performance Standards for Antimicrobial Susceptibility Testing (30th Edition). CLSI Clinical and Laboratory Standards Institute; 2015.

6. Liu C, Bayer A, Cosgrove SE, Daum RS, Fridkin SK, Gorwitz RJ, Kaplan SL, Karchmer AW, Levine DP, Murray BE, M JR, Talan DA, Chambers HF, Infectious Diseases Society of A. 2011. Clinical practice guidelines by the infectious diseases society of america for the treatment of methicillin-resistant Staphylococcus aureus infections in adults and children. Clin Infect Dis 52:e18–55.

7. Zuniga E, Roberti Benites N, da Hora AS, Mello PL, Laes MA, De Oliveira LAR, Brandao PE, Silva SOS, Taniwaki SA, Melville PA. 2020. Expression of genes encoding resistance in Staphylococcus spp. isolated from bovine subclinical mastitis in Brazil. J Infect Dev Ctries 14:772–780.

8. Geriak M, Haddad F, Rizvi K, Rose W, Kullar R, LaPlante K, Yu M, Vasina L, Ouellette K, Zervos M, Nizet V, Sakoulas G. 2019. Clinical Data on Daptomycin plus Ceftaroline versus Standard of Care Monotherapy in the Treatment of Methicillin-Resistant Staphylococcus aureus Bacteremia. Antimicrob Agents Chemother 63.

9. Tong SYC, Lye DC, Yahav D, Sud A, Robinson JO, Nelson J, Archuleta S, Roberts MA, Cass A, Paterson DL, Foo H, Paul M, Guy SD, Tramontana AR, Walls GB, McBride S, Bak N, Ghosh N, Rogers BA, Ralph AP, Davies J, Ferguson PE, Dotel R, McKew GL, Gray TJ, Holmes NE, Smith S, Warner MS, Kalimuddin S, Young BE, Runnegar N, Andresen DN, Anagnostou NA, Johnson SA, Chatfield MD, Cheng AC, Fowler VG, Jr., Howden BP, Meagher N, Price DJ, van Hal SJ, O’Sullivan MVN, Davis JS, Australasian Society for Infectious Diseases Clinical Research N. 2020. Effect of Vancomycin or Daptomycin With vs Without an Antistaphylococcal beta-Lactam on Mortality, Bacteremia, Relapse, or Treatment Failure in Patients With MRSA Bacteremia: A Randomized Clinical Trial. JAMA 323:527–537.

10. Hughes D, Andersson DI. 2017. Environmental and genetic modulation of the phenotypic expression of antibiotic resistance. FEMS Microbiol Rev 41:374–391.

11. Ersoy SC, Heithoff DM, Barnes Lt, Tripp GK, House JK, Marth JD, Smith JW, Mahan MJ. 2017. Correcting a Fundamental Flaw in the Paradigm for Antimicrobial Susceptibility Testing. EBioMedicine 20:173–181.

12. Radlinski L, Conlon BP. 2018. Antibiotic efficacy in the complex infection environment. Curr Opin Microbiol 42:19–24.

13. Knott S, Curry D, Zhao N, Metgud P, Dastgheyb SS, Purtill C, Harwood M, Chen AF, Schaer TP, Otto M, Hickok NJ. 2021. Staphylococcus aureus Floating Biofilm Formation and Phenotype in Synovial Fluid Depends on Albumin, Fibrinogen, and Hyaluronic Acid. Front Microbiol 12:655873.

14. Kuehne SA, Dempster AW, Collery MM, Joshi N, Jowett J, Kelly ML, Cave R, Longshaw CM, Minton NP. 2018. Characterization of the impact of rpoB mutations on the in vitro and in vivo competitive fitness of Clostridium difficile and susceptibility to fidaxomicin. J Antimicrob Chemother 73:973–980.

15. Cohen Y, Hershberg R. 2022. Rapid Adaptation Often Occurs through Mutations to the Most Highly Conserved Positions of the RNA Polymerase Core Enzyme. Genome Biol Evol 14.

16. Panchal VV, Griffiths C, Mosaei H, Bilyk B, Sutton JAF, Carnell OT, Hornby DP, Green J, Hobbs JK, Kelley WL, Zenkin N, Foster SJ. 2020. Evolving MRSA: High-level beta-lactam resistance in Staphylococcus aureus is associated with RNA Polymerase alterations and fine tuning of gene expression. PLoS Pathog 16:e1008672.

17. Aiba Y, Katayama Y, Hishinuma T, Murakami-Kuroda H, Cui L, Hiramatsu K. 2013. Mutation of RNA polymerase beta-subunit gene promotes heterogeneous-to-homogeneous conversion of beta-lactam resistance in methicillin-resistant Staphylococcus aureus. Antimicrob Agents Chemother 57:4861–71.

18. Patel Y, Soni V, Rhee KY, Helmann JD. 2023. Mutations in rpoB That Confer Rifampicin Resistance Can Alter Levels of Peptidoglycan Precursors and Affect beta-Lactam Susceptibility. mBio 14:e0316822.

19. Kato F, Sugai M. 2011. A simple method of markerless gene deletion in Staphylococcus aureus. J Microbiol Methods 87:76–81.

20. Mistretta N, Brossaud M, Telles F, Sanchez V, Talaga P, Rokbi B. 2019. Glycosylation of Staphylococcus aureus cell wall teichoic acid is influenced by environmental conditions. Sci Rep 9:3212.

21. Gerlach D, Guo Y, De Castro C, Kim SH, Schlatterer K, Xu FF, Pereira C, Seeberger PH, Ali S, Codee J, Sirisarn W, Schulte B, Wolz C, Larsen J, Molinaro A, Lee BL, Xia G, Stehle T, Peschel A. 2018. Methicillin-resistant Staphylococcus aureus alters cell wall glycosylation to evade immunity. Nature 563:705–709.

22. Schlag M, Biswas R, Krismer B, Kohler T, Zoll S, Yu W, Schwarz H, Peschel A, Gotz F. 2010. Role of staphylococcal wall teichoic acid in targeting the major autolysin Atl. Mol Microbiol 75:864–73.

23. Hort M, Bertsche U, Nozinovic S, Dietrich A, Schrotter AS, Mildenberger L, Axtmann K, Berscheid A, Bierbaum G. 2021. The Role of beta-Glycosylated Wall Teichoic Acids in the Reduction of Vancomycin Susceptibility in Vancomycin-Intermediate Staphylococcus aureus. Microbiol Spectr 9:e0052821.

24. Xia G, Maier L, Sanchez-Carballo P, Li M, Otto M, Holst O, Peschel A. 2010. Glycosylation of wall teichoic acid in Staphylococcus aureus by TarM. J Biol Chem 285:13405–15.

25. Miyachiro MM, Contreras-Martel C, Dessen A. 2019. Penicillin-Binding Proteins (PBPs) and Bacterial Cell Wall Elongation Complexes. Subcell Biochem 93:273–289.

26. Goldstein BP. 2014. Resistance to rifampicin: a review. J Antibiot (Tokyo) 67:625–30.

27. Li MC, Lu J, Lu Y, Xiao TY, Liu HC, Lin SQ, Xu D, Li GL, Zhao XQ, Liu ZG, Zhao LL, Wan KL. 2021. rpoB Mutations and Effects on Rifampin Resistance in Mycobacterium tuberculosis. Infect Drug Resist 14:4119–4128.

28. Guo Y, Wang B, Rao L, Wang X, Zhao H, Li M, Yu F. 2021. Molecular Characteristics of Rifampin-Sensitive and -Resistant Isolates and Characteristics of rpoB Gene Mutations in Methicillin-Resistant Staphylococcus aureus. Infect Drug Resist 14:4591–4600.

29. Wielders CL, Fluit AC, Brisse S, Verhoef J, Schmitz FJ. 2002. mecA gene is widely disseminated in Staphylococcus aureus population. J Clin Microbiol 40:3970–5.

30. Campbell EA, Korzheva N, Mustaev A, Murakami K, Nair S, Goldfarb A, Darst SA. 2001. Structural mechanism for rifampicin inhibition of bacterial rna polymerase. Cell 104:901–12.

31. Kristich CJ, Little JL. 2012. Mutations in the beta subunit of RNA polymerase alter intrinsic cephalosporin resistance in Enterococci. Antimicrob Agents Chemother 56:2022–7.

32. Kho K, Meredith TC. 2018. Extraction and Analysis of Bacterial Teichoic Acids. Bio Protoc 8:e3078.

33. Rajagopal M, Walker S. 2017. Envelope Structures of Gram-Positive Bacteria. Curr Top Microbiol Immunol 404:1–44.

34. Rohde M. 2019. The Gram-Positive Bacterial Cell Wall. Microbiol Spectr 7.

35. Baur S, Rautenberg M, Faulstich M, Grau T, Severin Y, Unger C, Hoffmann WH, Rudel T, Autenrieth IB, Weidenmaier C. 2014. A nasal epithelial receptor for Staphylococcus aureus WTA governs adhesion to epithelial cells and modulates nasal colonization. PLoS Pathog 10:e1004089.

36. Percy MG, Grundling A. 2014. Lipoteichoic acid synthesis and function in gram-positive bacteria. Annu Rev Microbiol 68:81–100.

37. Wu X, Han J, Gong G, Koffas MAG, Zha J. 2021. Wall teichoic acids: physiology and applications. FEMS Microbiol Rev 45.

38. Neuhaus FC, Baddiley J. 2003. A continuum of anionic charge: structures and functions of D-alanyl-teichoic acids in gram-positive bacteria. Microbiol Mol Biol Rev 67:686–723.

